# APOBEC3B mRNA Expression in Breast Cancer Correlates with Genomic Mutational Signatures

**DOI:** 10.64898/2026.08.20.745899

**Authors:** Joel Pardo, Nuri A. Temiz, Douglas Yee

## Abstract

Despite advances in screening and treatment, breast cancer remains a leading cause of cancer-related mortality. APOBEC enzymes, particularly APOBEC3B (A3B), are upregulated in many cancers, contributing to a characteristic C-to-T mutational signature found in 30-50% of breast cancers. However, the relationship between A3B mutational signatures and A3B expression across subtypes, and the resulting potential biologic consequences, have not been fully defined. Using TCGA and ICGC datasets, we analyzed DNA and RNA expression data to assess the relationship between *A3B* mRNA expression and APOBEC enrichment scores. Pathway enrichment analyses (KEGG, GO, Reactome) were performed to identify biological processes associated with high A3B expression, specifically stratifying by breast cancer intrinsic subtypes (HR+/HER2−, HR+/HER2+, HR−/HER2+, and TNBC). Over 64% of tumors with enriched A3B mutational genomic signatures demonstrated above-median *A3B* mRNA expression (p < 0.001). High A3B-expressing tumors exhibited specific alterations in drug metabolism pathways. Notably, we observed reduced expression of *CYP2D6* and *CYP3A* isoforms which is required for the conversion of tamoxifen to its active metabolites. Conversely, genes involved in pyrimidine metabolism, including *IMPDH1, NME1, TK1*, and *DPYS*, were downregulated in high A3B tumors. Elevated A3B expression correlates with mutational signatures and may contribute to impaired tamoxifen activation and endocrine resistance, while concurrently creating metabolic vulnerabilities to pyrimidine-based chemotherapies. Targeting A3B or exploiting these metabolic dependencies may improve therapeutic response in selected patient subsets.

## Introduction

Breast cancer remains a leading cause of women’s cancer-related mortality despite advances in early detection and targeted therapies [1, 2]. The complex molecular landscape of breast cancer continues to drive research into novel diagnostic and therapeutic strategies. A growing body of evidence suggests that endogenous mutagenic processes contribute to tumor evolution and therapeutic resistance [3].

Among these processes, the APOBEC (apolipoprotein B mRNA-editing enzyme catalytic polypeptide-like) family of cytosine deaminases has emerged as a key driver of genomic instability in multiple cancer types, including breast cancer [4]. These enzymes, primarily tasked with antiviral defense, catalyze cytosine-to-uracil deamination, generating C-to-T and C-to-G mutations that can accumulate over time [5]. *APOBEC3B* (*A3B*), in particular, is frequently overexpressed in breast cancer and has been implicated in tumor progression, immune evasion, and drug resistance [6-8]. While the presence of an A3B-associated mutational signature has been identified in a substantial subset of breast cancers, its relationship with *A3B* mRNA expression and downstream biological consequences remains incompletely understood [9].

Recent studies have highlighted the potential impact of A3B expression on treatment response, particularly in relation to endocrine and chemotherapeutic agents [7, 10, 11]. High A3B expression has been associated with increased tumor mutational burden and greater intratumoral heterogeneity [12]. Notably, prior research suggests that elevated A3B levels may impair the therapeutic efficacy of tamoxifen, a selective estrogen receptor modulator widely used in hormone receptor-positive breast cancer [7].

The molecular pathways influencing A3B expression and activity, particularly those involved in drug metabolism, are areas of growing interest. While prior studies have established a correlation between A3B activity and increased tumor mutational burden, the precise relationship between *A3B* mRNA expression and the accumulation of A3B-driven mutational signatures across different clinical subtypes remains underexplored [13, 14]. Despite recent insights, a comprehensive understanding of how A3B expression alters the tumor’s metabolic dependencies and contributes to therapy resistance is incomplete. Elucidating these relationships is critical, as A3B-driven mutagenesis could serve not merely as a biomarker of genomic instability, but as a functional driver of specific therapeutic vulnerabilities.

Our findings highlight a robust relationship between *A3B* mRNA expression and APOBEC mutational enrichment. Furthermore, exploration of the pathways associated with A3B expression revealed specific alterations in metabolism pathways of drugs frequently used as breast cancer treatment, including tamoxifen and 5-fluorouracil. Understanding these pathways offers new insights into the molecular mechanisms driving breast cancer progression and may guide the selection of therapies for patients with A3B-enriched tumors [15].

## Methods

### Data acquisition and preprocessing

Genomic and transcriptomic data for breast cancer patients were obtained from The Cancer Genome Atlas (TCGA) and the International Cancer Genome Consortium (ICGC) databases. TCGA data included DNA sequencing (whole-exome) and RNA sequencing (RNA-Seq) data for primary breast tumors, while ICGC data provided RNA sequencing information for additional breast cancer cohorts. Clinical metadata, including breast cancer subtypes (HR+/HER2−, HR+/HER2+, HR−/HER2+, and TNBC), were extracted from these databases (Fig 1).

**Fig 1.**
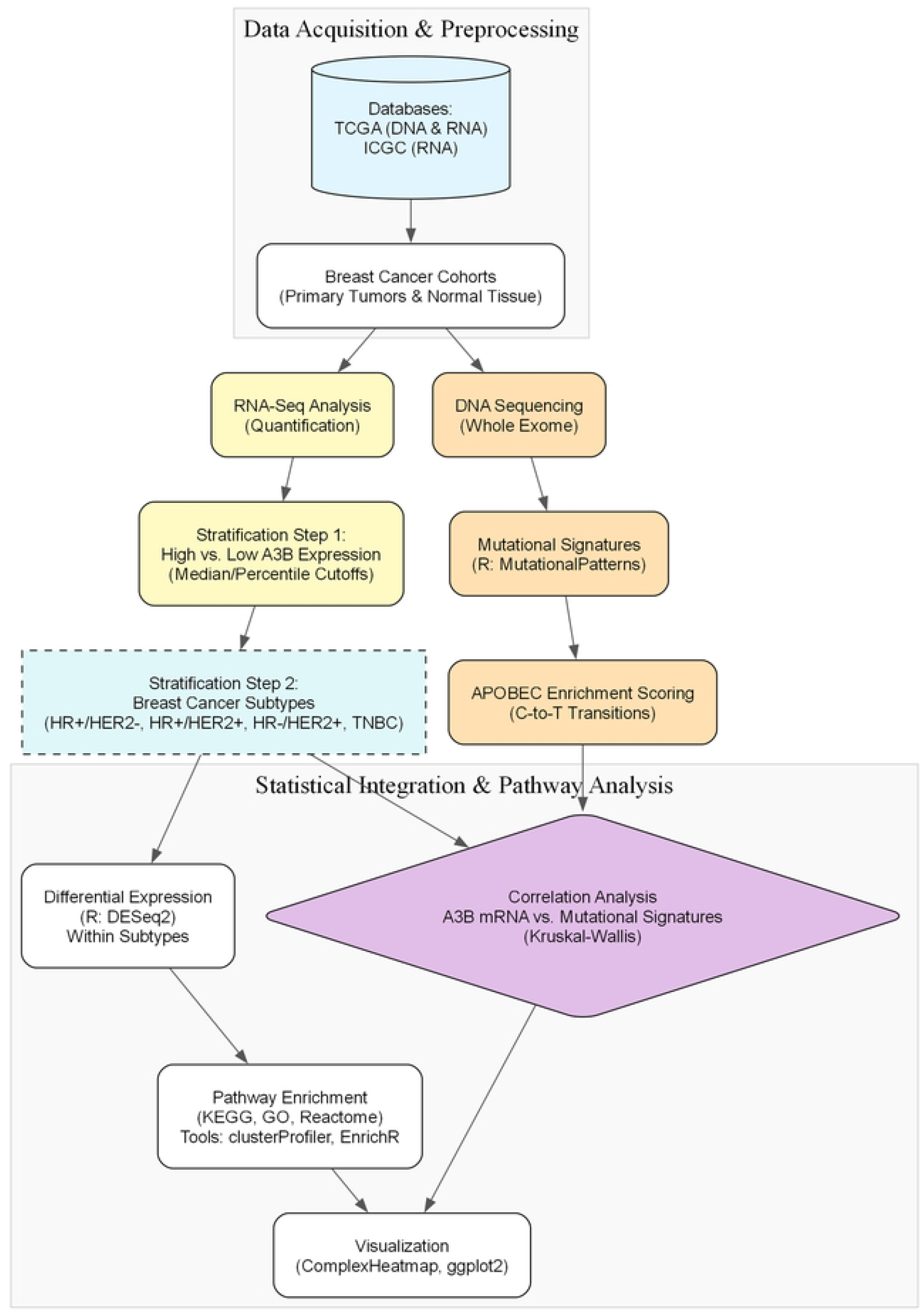
Analytical workflow for data acquisition, processing, and pathway analysis. Flowchart illustrating the bioinformatic methodology used in the study. Genomic and transcriptomic data were acquired from primary breast tumor and normal tissue cohorts within the TCGA and ICGC databases. RNA-Seq data was utilized to quantify expression and stratify samples into high versus low APOBEC3B (A3B) expression groups using 33^rd^ percentile cutoffs. Concurrently, whole-exome DNA sequencing data was processed via the MutationalPatterns R package to derive APOBEC enrichment scores based on C-to-T transitions. Statistical integration involved Kruskal-Wallis correlation analysis between A3B mRNA levels and mutational signatures, followed by differential expression analysis (DESeq2) stratified by intrinsic breast cancer subtypes (HR+/HER2−, HR+/HER2+, HR−/HER2+, TNBC). Subsequent pathway enrichment (KEGG, GO, Reactome) was performed using clusterProfiler and EnrichR, with final visualizations generated via ComplexHeatmap and ggplot2

### A3B expression and mutational signature analysis

*APOBEC3B* (*A3B*) expression levels were quantified using normalized RNA-Seq data. Tumors were stratified into high and low A3B expression groups based on the median A3B expression level across the entire cohort. Mutational signatures were identified using the MutationalPatterns package in R, focusing on the APOBEC-related C-to-T transitions [31].

### Pathway enrichment analysis

Pathway enrichment analysis was conducted using KEGG, Gene Ontology (GO), and Reactome databases. Differentially expressed genes between high and low A3B-expressing tumors were identified using the DESeq2 package in R [32]. Additional differential comparison was performed using a non-parametric t-test with Benjamini-Hochberg correction for multiple testing [33]. Significantly altered genes were then analyzed for pathway enrichment using the clusterProfiler and EnrichR package [34]. Heatmaps of pathway enrichment scores were generated using the ComplexHeatmap package [35]. KEGG terms were retrieved using org.Hs.eg.db, GO terms were obtained using the Gene Ontology knowledgebase, and Reactome pathway annotations were derived from the Reactome knowledgebase [35-38].

### Breast cancer subtype analysis

To investigate the role of A3B across different breast cancer subtypes, we stratified the dataset into hormone receptor-positive/HER2− positive (HR+/HER2+), hormone receptor-positive/HER2− negative (HR+/HER2−), hormone receptor-negative/HER2− positive (HR−/HER2+), and triple-negative breast cancer (TNBC) subtypes based on clinical annotations from TCGA and ICGC. A3B expression levels and mutational signature enrichment were analyzed within each subtype, and pathway enrichment analysis was performed as described above. The distribution of A3B expression across subtypes was visualized using ggplot2, while pathway enrichment differences were displayed using heatmaps and ggplot2.

### Statistical analysis

Statistical analyses were performed using R (version 4.4.2). Data visualization was carried out using ggplot2 [39] and ComplexHeatmap [35] libraries in R. Differences were considered statistically significant at p < 0.05.

## Results

### High A3B expression correlates with APOBEC mutational signature enrichment

To examine the relationship between APOBEC mutational signature enrichment and A3B expression levels in breast cancer, we analyzed DNA and RNA expression data from TCGA-BRCA (n = 1097 tumor samples and n = 113 adjacent normal breast tissue samples). We defined “high A3B” as samples with A3B mRNA expression above the 33rd percentile after normalizing APOBEC3B expression by TATA-box binding protein (TBP, a house keeping gene). This threshold corresponded to above the 80^th^ percentile in A3B expression for normal breast tissue (Supplemental 1).

**Supplemental Figure 1. Establishment of APOBEC3B expression thresholds using normal tissue controls. (A)** Density plot comparing raw log2(Expression + 1) values between primary tumors (n = 1093) and solid tissue normal samples (n = 112). The dashed red line marks the 33rd percentile threshold (6.74) for tumors. **(B)** Density plot of the housekeeping gene TBP across primary tumor and normal tissue samples. **(C)** Waterfall plot depicting the log2 expression of APOBEC3B normalized by TBP, grouped by breast cancer subtype. Dashed lines represent subtype means, while the solid black line denotes the 33rd percentile cut-off (−1.2). Shape indicates significant APOBEC enrichment.

We observed a significant association between A3B expression and APOBEC mutational signature enrichment (Fig 2A and 2B). Tumors with high A3B expression displayed higher APOBEC enrichment scores compared to those with low expression (p < 0.05, Kruskal-Wallis test) (Fig 2B). Samples with an adjusted p-value < 0.05 for the enrichment score were classified as having high APOBEC mutational activity (Fig 2A). The distribution of A3B expression, visualized via waterfall plot, confirms that the highest A3B expression levels are dominated by specific aggressive subtypes. Notably, the upper quantile of expression is populated densely by HR−/HER2+ samples (orange triangles) and TNBC samples (light blue circles), whereas HR+ samples are more evenly distributed across the expression spectrum (Fig 2A, S1 Fig). These findings indicate that elevated A3B expression is a strong predictor of increased APOBEC-driven mutagenesis, particularly in HER2− enriched and triple-negative subtypes.

**Fig 2.**
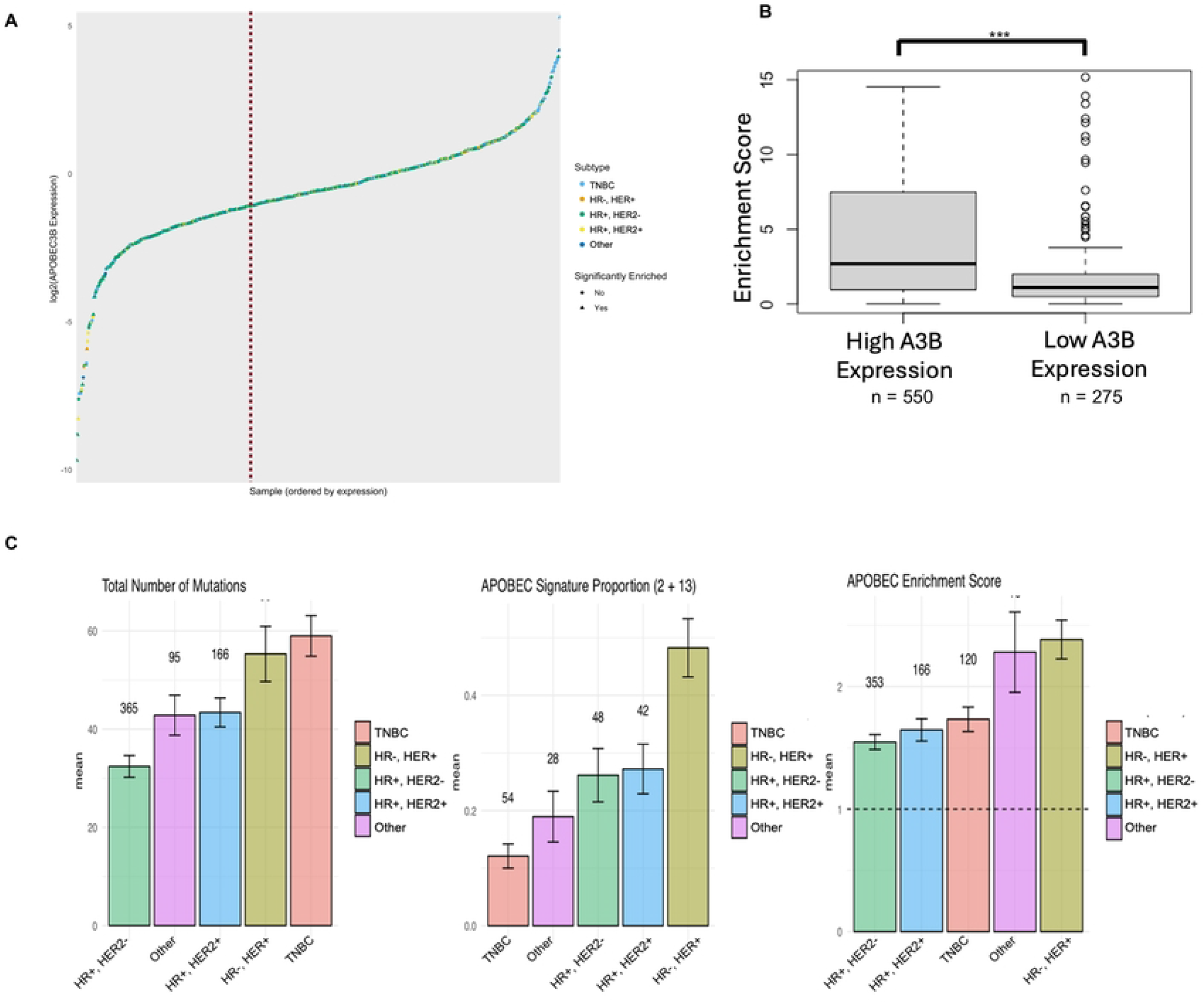
APOBEC3B expression and its association with mutational signatures across breast cancer subtypes. **(A)** Waterfall plot showing the distribution of log2(APOBEC3B Expression) across ordered samples. Individual data points are colored by intrinsic breast cancer subtype (TNBC, HR−/HER2+, HR+/HER2−, HR+/HER2+, Other), and shapes (circles vs. triangles) denote the absence or presence of significant APOBEC enrichment, respectively. A dashed vertical red line indicates the expression threshold separating high and low A3B groups. **(B)** Boxplot comparing APOBEC enrichment scores between High A3B expression (n = 550) and Low A3B expression (n = 275) groups, demonstrating significantly higher enrichment in the high-expression cohort. **(C)** Bar charts detailing the mean total number of mutations, APOBEC signature proportion (signatures 2 + 13), and APOBEC enrichment score, stratified by breast cancer subtype. Error bars represent the standard error of the mean.

### Subtype-Specific Patterns of APOBEC Activity

We next evaluated how these patterns differed across intrinsic breast cancer subtypes. As expected, TNBC exhibited the highest overall mutational burden, followed by HR−/HER2+, HR+/HER2+, and HR+/HER2− tumors. However, the distribution of APOBEC-associated mutations differed from the total mutation load.

HR−/HER2+ tumors displayed the highest APOBEC signature 2/13 mutation counts, whereas TNBC, despite having the highest overall mutation burden, showed comparatively fewer APOBEC-specific mutations (Fig 2C). When considering the APOBEC enrichment score, which accounts for the clustered context of APOBEC signature mutations, TNBC rose to the second highest score, just below HR−/HER2+. A3B mRNA expression mirrored this enrichment pattern: HR−/HER2+ tumors had the highest A3B expression, followed by TNBC (Fig 2C).

This division between total mutations and APOBEC-specific signatures implies that while TNBC is genomically unstable due to multiple overlapping processes, HR−/HER2+ tumors may be uniquely and disproportionately dependent on APOBEC-mediated mutagenesis.

### Altered Drug Metabolism Pathways in A3B-High Tumors

To clarify biological processes associated with A3B expression, we performed pathway enrichment analyses (KEGG, GO, Reactome). Across TCGA samples, tumors with high A3B expression showed significant enrichment in pathways related to drug metabolism, immune signaling, and cell cycle regulation (Fig 3).

**Fig 3.**
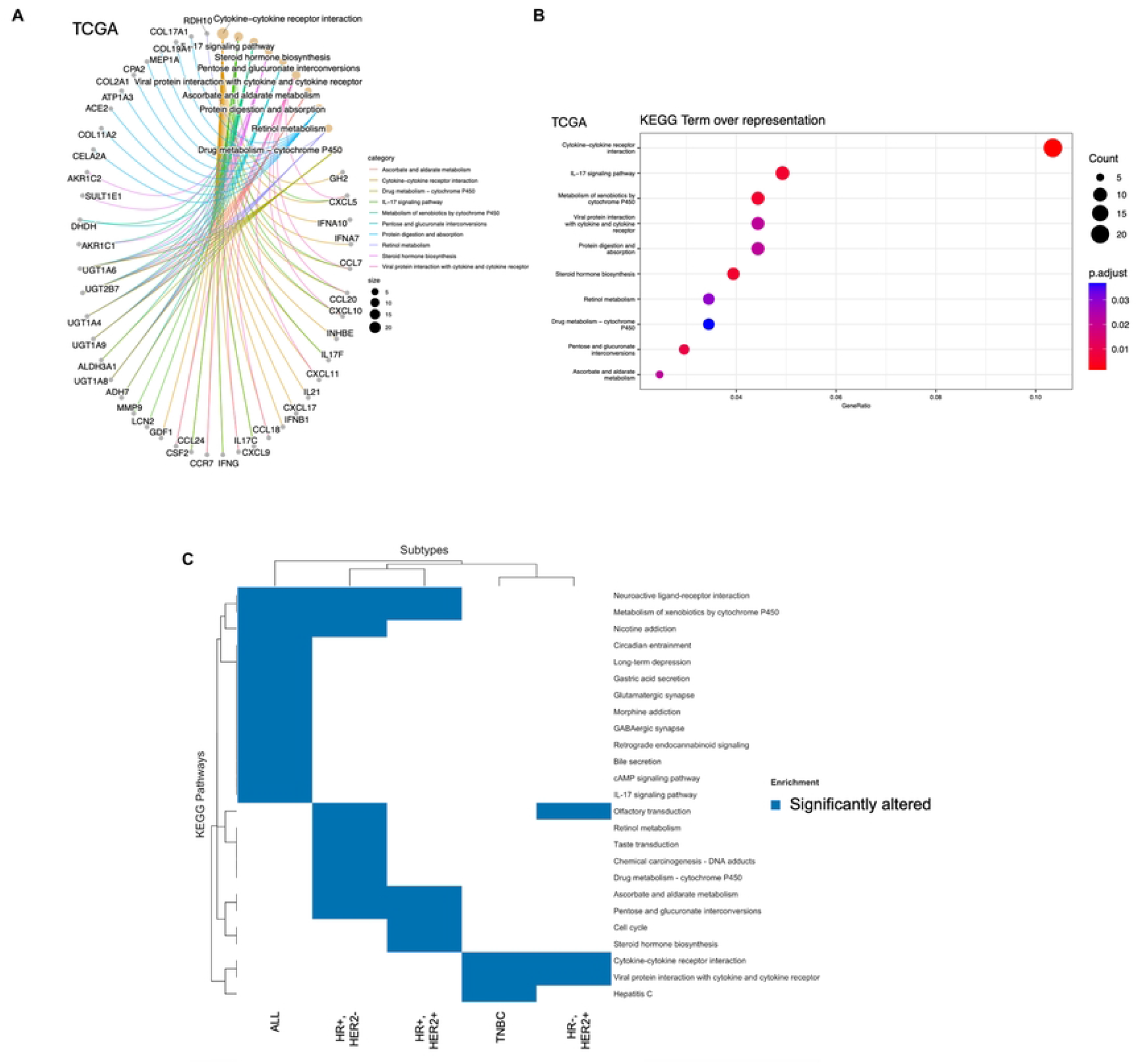
Pathway enrichment and altered drug metabolism networks in A3B-high breast tumors (TCGA Cohort). **(A)** Network plot illustrating the connections between significantly altered genes and over-represented KEGG pathways, including drug metabolism (cytochrome P450), retinol metabolism, and cytokine-cytokine receptor interactions. Node size reflects the gene count, and line colors correspond to specific pathway categories. **(B)** Dot plot displaying KEGG term over-representation analysis. The x-axis represents the GeneRatio, dot size indicates the count of associated genes, and color gradients reflect adjusted p-values. **(C)** Heatmap showing hierarchical clustering of significantly altered KEGG pathways across distinct breast cancer subtypes (ALL, HR+/HER2−, HR+/HER2+, TNBC, HR−/HER2+). Blue blocks indicate significant enrichment for the corresponding pathway within that subtype.

Within drug metabolism pathways, A3B-high tumors demonstrated reduced expression of key cytochrome P450 enzymes required for tamoxifen activation, including *CYP2D6* and *CYP3A4/3A5*, particularly within HR+/HER2− and HR+/HER2+ subtypes. This pattern suggests that elevated A3B expression may impair the conversion of tamoxifen to its active metabolites.

In contrast, changes affecting cyclophosphamide metabolism (e.g., reduced *CYP2B6*) and broader xenobiotic metabolism pathways were observed across multiple subtypes, including TNBC. Additionally, many genes contributing to these pathways overlapped with immune activation networks (including *CCL5, CCL20, IFNG*), which were enriched in TNBC and HR−/HER2+ tumors but less prominent in HR+/HER2− tumors (Fig 3).

### Metabolic Vulnerabilities and Therapeutic Implications

In A3B-high tumors, we observed downregulation (green nodes) of *CYP2D6*, the rate-limiting enzyme required to convert tamoxifen into its active metabolite, endoxifen. In contrast, *CYP1A1* (labeled 1.14.13.8) showed upregulation (red node). The specific suppression of *CYP2D6* suggests a mechanism for intrinsic tamoxifen resistance in A3B-high HR+ patients that is distinct from genomic polymorphisms (Fig 4A).

**Fig 4.**
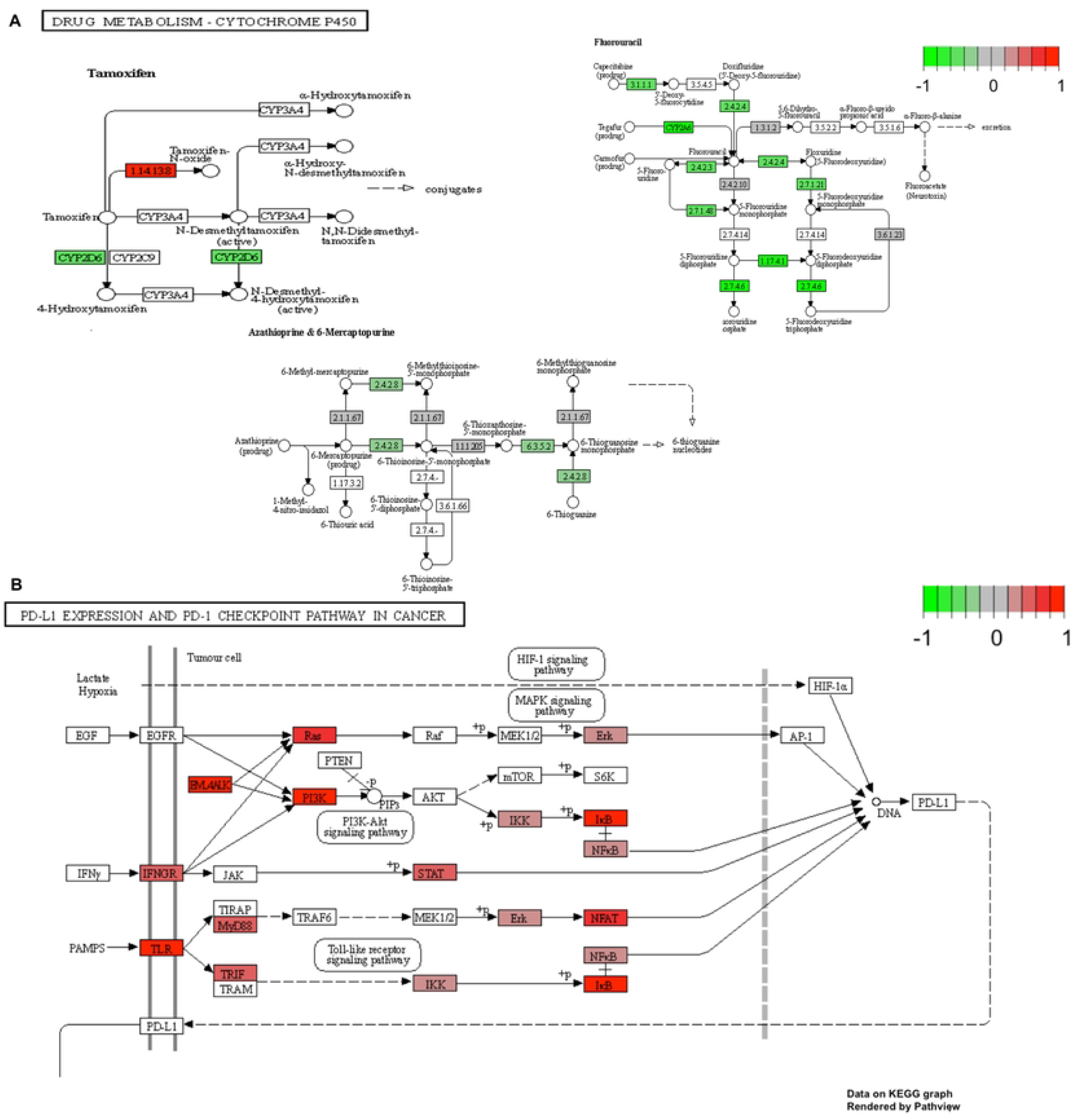
Metabolic vulnerabilities and immune-related pathway alterations associated with high A3B expression. KEGG pathway maps rendered by Pathview detailing specific gene expression changes, where red nodes indicate upregulation, and green nodes indicate downregulation. **(A)** The Drug Metabolism - Cytochrome P450 pathway map centered on tamoxifen metabolism highlights the downregulation of CYP2D6 (green) and upregulation of 1.14.13.8/CYP1A1 (red). A supplementary map for fluorouracil metabolism shows the downregulation of key catabolic enzymes, including 3.5.2.2/DPYS. **(B)** The PD-L1 expression and PD-1 checkpoint pathway demonstrates the upregulation of key inflammatory mediators, including STAT, NFκB, PI3K, and TLR (red nodes).

Genes involved in the catabolism of pyrimidines were also modulated. The pathway map for Fluorouracil metabolism shows downregulation (green nodes) of enzymes such as *DPYS* (dihydropyrimidinase, 3.5.2.2) and *UPB1* (beta-ureidopropionase) (Fig 4A). Reduced catabolic activity in this pathway theoretically increases the intracellular half-life of active 5-FU metabolites, suggesting hypersensitivity to fluoropyrimidine-based therapy.

In parallel, immune-related pathways were significantly upregulated. We observed upregulation (red nodes) of key inflammatory mediators including *STAT, NFkB, PI3K*, and *TLR* within the PD-L1 expression and PD-1 checkpoint pathway (Fig 4B). This was corroborated by ICGC data, where “PD-L1 expression” and “Natural killer cell mediated cytotoxicity” were top enriched terms, particularly in TNBC samples (S2 Fig).

**Supplemental Figure 2. Validation of pathway enrichment in the ICGC cohort and correlations with APOBEC3A and tumor mutational burden. (A)** Dot plot displaying KEGG term over-representation for the ICGC dataset, highlighting the enrichment of immune-related pathways such as natural killer cell mediated cytotoxicity and cytokine-cytokine receptor interaction. **(B)** Network plot for the ICGC cohort mapping altered genes to their respective KEGG pathways. **(C)** Heatmap of KEGG pathway enrichment across subtypes in the ICGC cohort, where yellow blocks signify significant alterations.

## Discussion

Our study provides insights into the role of *APOBEC3B* (*A3B*) in breast cancer, particularly its relationship to mutational processes, pathway reprogramming, and treatment response. Using TCGA dataset, we confirmed that higher A3B expression is significantly associated with tumors enriched for APOBEC mutational signatures. The boxplot analysis (Fig 2B) reinforces that high A3B expression serves as a robust proxy for active APOBEC mutagenesis, with enrichment scores in the high-expression group consistently exceeding those in the low-expression group.

A3B expression and APOBEC enrichment varied significantly across subtypes. Although TNBC had the highest total mutation burden, HR−/HER2+ tumors demonstrated the strongest APOBEC signature burden (Fig 2C). This distinction reinforces that A3B expression correlates more closely with the APOBEC signature enrichment score than with total mutational burden. A targeted analysis comparing A3B expression and TMB in TCGA is included in S3 Fig.

**Supplemental Figure 3. APOBEC3B expression compared to TMB, APOBEC3A expression, and APOBEC3A mutational patterns. (A)** Spearman correlation between APOBEC3B (x-axis) and Tumor Mutational Burden (TMB, y-axis). **(B)** Scatter plot revealing a positive correlation (R^2^ = 0.289, Spearman ρ = 0.573) between log2(APOBEC3B + 1) and log2(APOBEC3A + 1) expression levels. **(C)** Violin plots illustrate the distribution of the log2-transformed ratio of APOBEC-associated mutations in the YTCA context relative to the RTCA context. Data is stratified by clinical breast cancer subtype. Jittered overlaid points represent individual tumor samples.

A practical challenge in studying APOBEC biology is distinguishing A3B from the closely related A3A due to high sequence homology. In our analysis, we found a moderate correlation between A3B and A3A expression (p-value = 0.573, Spearman), yet A3B expression levels were generally higher and more pervasive across samples (S3 Fig). This supports the utility of A3B as the primary biomarker for this mutational process in breast cancer.

Crucially, pathway analyses revealed that A3B-high tumors exhibit altered drug metabolism profiles. In HR+ subtypes, high A3B expression was associated with reduced expression of *CYP2D6*, the enzyme critical for tamoxifen bioactivation. This downregulation (visualized as green nodes in Fig 4A) provides a genotype-independent mechanism for endocrine resistance. Conversely, the downregulation of pyrimidine catabolism genes like *DPYS* in A3B-high tumors (Fig 4B) may increase intracellular retention of 5-fluorouracil metabolites, indicating potential therapeutic sensitivity to fluoropyrimidines.

Furthermore, the upregulation of PD-L1 checkpoint pathway components (*STAT, NFkB*) in A3B-high tumors, particularly in TNBC, suggests that these tumors may be primed for immune checkpoint blockade, consistent with their higher mutational load and neoantigen potential.

This study is limited by its correlative nature. While genome-wide sequencing remains the gold standard for detecting APOBEC patterns, A3B expression could offer a practical surrogate that is easier to measure clinically.

## Conclusions

A3B expression marks a subset of breast cancers with distinct mutational processes and drug metabolism programs. Our findings suggest that elevated A3B expression may contribute to impaired tamoxifen activation and endocrine resistance, while concurrently creating metabolic vulnerabilities to pyrimidine-directed therapies. Integrating A3B expression into clinical decision-making may ultimately help guide endocrine therapy selection and identify patients more likely to benefit from fluoropyrimidine-based treatment.

## Declarations

### Ethics approval and consent to participate

Not applicable for this study as it utilized publicly available data (TCGA and ICGC).

### Availability of data and materials

The datasets generated and/or analyzed during the current study are available in the TCGA repository (https://portal.gdc.cancer.gov/) and ICGC data portal (https://dcc.icgc.org/).

### Authors’ contributions

JP performed bioinformatic analysis, analyzed the data and wrote the manuscript. NAT supervised the bioinformatic analysis, analyzed the data, and edited the manuscript. DY supervised the study and edited the manuscript. All authors read and approved the final manuscript.

### Consent for publication

All authors have consented for this work to be considered for publication.

## Acknowledgments

Not applicable.

